# AIM: A Network Model of Attention in Auditory Cortex

**DOI:** 10.1101/2020.12.10.419762

**Authors:** Kenny F. Chou, Kamal Sen

## Abstract

Attentional modulation of cortical networks is critical for the cognitive flexibility required to process complex scenes. Current theoretical frameworks for attention are based almost exclusively on studies in visual cortex, where attentional effects are typically modest and excitatory. In contrast, attentional effects in auditory cortex can be large and suppressive. A theoretical framework for explaining attentional effects in auditory cortex is lacking, preventing a broader understanding of cortical mechanisms underlying attention. Here, we present a cortical network model of attention in primary auditory cortex (A1). A key mechanism in our network is attentional inhibitory modulation (AIM) of cortical inhibitory neurons. In this mechanism, top-down inhibitory neurons disinhibit bottom-up cortical circuits, a prominent circuit motif observed in sensory cortex. Our results reveal that the same underlying mechanisms in the AIM network can explain diverse attentional effects on both spatial and frequency tuning in A1. We find that a dominant effect of disinhibition on cortical tuning is suppressive, consistent with experimental observations. Functionally, the AIM network may play a key role in solving the cocktail party problem. We demonstrate how attention can guide the AIM network to monitor an acoustic scene, select a specific target, or switch to a different target, providing flexible outputs for solving the cocktail party problem.

## Introduction

A hallmark of cortical processing is the capacity for generating flexible behaviors in a context-dependent manner. A striking example of a problem that requires such cognitive flexibility is the cocktail party problem, where a listener can selectively listen to a speaker amongst other speakers (McDermott, 2009). Listening in such settings can be highly flexible, depending on the goal of the listener. For example, a listener can *monitor* the entire auditory scene, *select* a particular target, or *switch* to another target. Recent theoretical and experimental studies have begun to propose model networks and cortical mechanisms for producing flexible behaviors (Vogels and Abbott, 2009; Letzkus et al., 2011; Pi et al., 2013; Zhang et al., 2014; Yang et al., 2016; Kuchibhotla et al., 2017; Wang and Yang, 2018); and top-down control of cortical circuits via attention is thought to be a critical component.

The influence of attention on cortical processing has been intensively investigated in vision, resulting in a prominent theoretical framework of attention (Desimone and Duncan, 1995; Reynolds and Heeger, 2009). In contrast, relatively little is known about attentional mechanisms in auditory cortex. After the early discovery of “attention units” in the auditory cortex (Hubel et al., 1959), there has recently been renewed interest on attentional effects in auditory cortex (Fritz et al., 2007b; Shinn-Cunningham, 2008; Otazu et al., 2009). In comparison to the effects of attention in primary visual cortex, which are relatively small and excitatory (Buffalo et al., 2010), attentional effects in primary auditory cortex (A1) can be much larger and suppressive (Fritz et al., 2003; Otazu et al., 2009; Lee and Middlebrooks, 2011). However, a theoretical framework for auditory cortical mechanisms underlying attention is lacking.

The responses of neurons in A1 can change rapidly when an animal is actively engaged in a task (Fritz et al., 2003; Otazu et al., 2009; Lee and Middlebrooks, 2011; Kuchibhotla et al., 2017). For example, cortical neurons with broad spatial tuning curves can sharpen tuning during attentive behavior (Lee and Middlebrooks, 2011); whereas the spectral temporal receptive fields (STRFs) of cortical neurons with narrow frequency tuning can display the emergence of entirely new excitatory regions (Fritz et al., 2003) or suppressive effects (Otazu et al., 2009). Cortical network mechanisms underlying such diverse attentional changes in tuning remain poorly understood. Changes in cortical tuning can also be driven by competing auditory stimuli in cocktail-party settings, even when an animal in anesthetized (Maddox et al., 2012; Middlebrooks and Bremen, 2013), suggesting the involvement of both bottom-up and top-down mechanisms (Fritz et al., 2007b; Bee and Micheyl, 2008; McDermott, 2009; Bronkhorst, 2015). Previous models of cortical processing underlying the cocktail party problem have largely focused on bottom-up mechanisms (Dong et al., 2016; Chou et al., 2019). Specific cortical circuit mechanisms underlying top-down attentional changes in cortical responses, and their functional role in solving the cocktail party problem, remain unclear (Kaya and Elhilali, 2017).

Here we propose a network model to explain how experimentally observed cortical response properties in A1 could arise from underlying network mechanisms, via the interplay between bottom-up and top-down processes. Central to our network model is attentional inhibitory modulation (AIM), i.e., attention-driven modulation of distinct populations of cortical inhibitory neurons. Specifically, this mechanism relies on disinhibition of bottom-up cortical circuits, mediated via top-down inhibitory neurons, a prominent motif observed in cortex (Letzkus et al., 2011; Pfeffer et al., 2013; Pi et al., 2013; Zhang et al., 2014; Kuchibhotla et al., 2017). We first use the AIM network to model diverse attentional changes in spatial and spectral tuning in auditory cortex (Fritz et al., 2003; Lee and Middlebrooks, 2011), and then illustrate its potential functional role in solving the cocktail party problem.

## Results

### The AIM Network

We began by focusing on spatial processing of multiple sound sources in auditory cortex, extending previous models of bottom-up processing (figure 1 A,B). The bottom-up network implements two key operations: integration and competition. Integration is mediated by broad convergence across spatial channels on the cortical neuron (C, Figure 1A); whereas competition is mediated by inhibition across spatial channels via I neurons (Figure 1B). These bottom-up mechanisms explain two key features observed experimentally in anesthetized or passive animals: broad spatial tuning of cortical neurons to single sounds; and sharpening of spatial tuning in the presence of multiple competing sounds (Maddox et al., 2012; Middlebrooks and Bremen, 2013; Dong et al., 2016; Chou et al., 2019).

**Figure 1:**
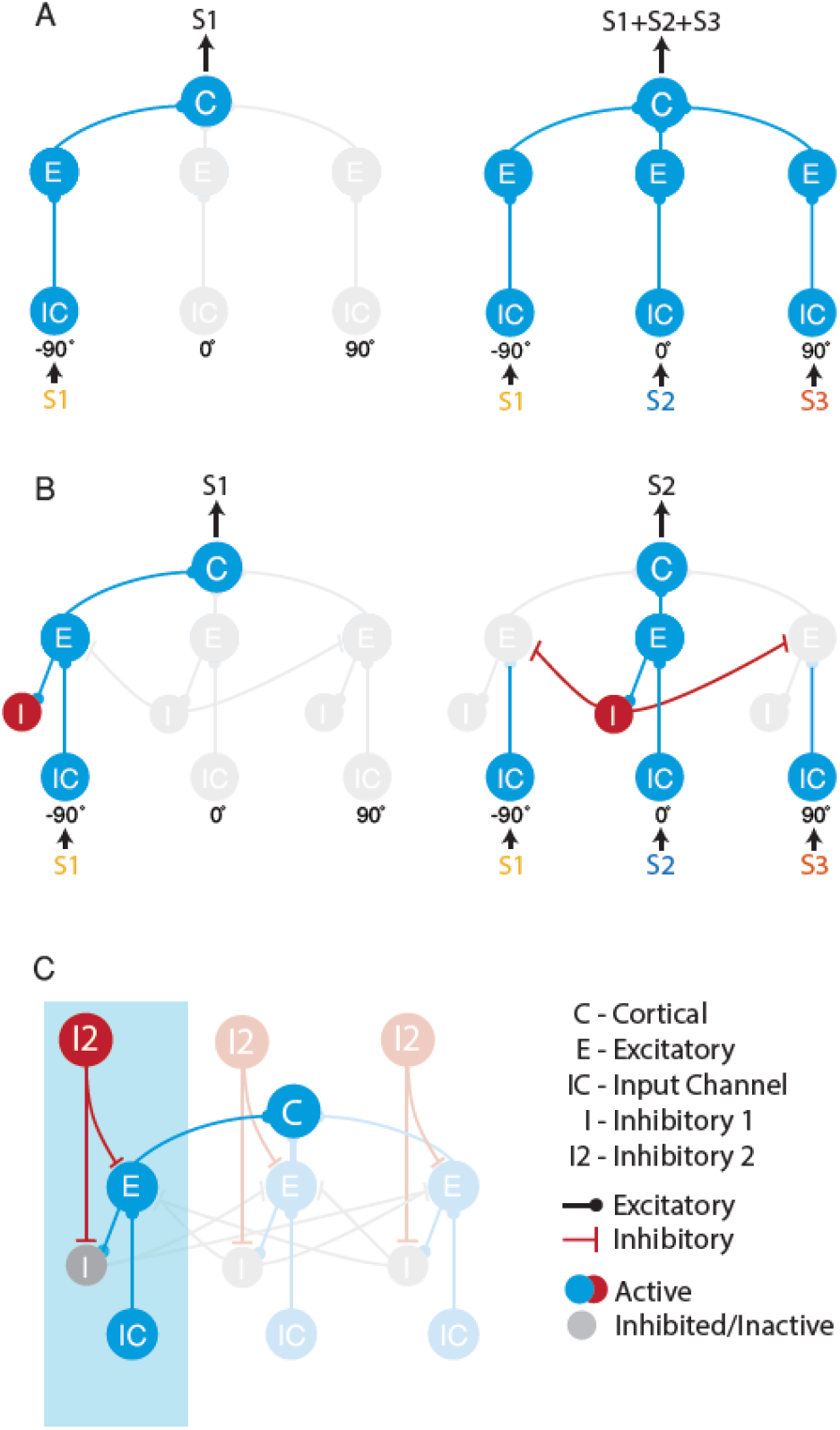
Sub-networks within the AIM network. (A) A convergence network. S1, S2, and S3 denote distinct audio stimuli 1, 2, and 3, which are placed virtually at −90°, 0°, and 90° azimuth, respectively. (B) A passive switching network realized with the addition of I neurons. (C) The AIM network, realized with the addition of I2 neurons, which modulates the I neurons. One spatial channel is highlighted in blue. Neurons in the same spatial channel processes the stimuli from a specific spatial location. Response of the cortical neuron represents the output of the network.

We extended the bottom-up network to model the effects of attention in the AIM network (Fig 1C). Previous studies have shown that bottom-up cortical representations can be modulated in the attentive state by distinct sub-types of inhibitory neurons. To model such attentional inhibitory modulation, we introduced an additional layer of inhibitory neurons (I2, Fig 1C). I2 neurons can control the spatial tuning of the cortical neuron in different attentive states, by modulating the activity of E and I neurons in the bottom-up network.

### Attentional Changes in Spatial Tuning

A previous experimental study in cat A1 demonstrated that the spatial tuning of cortical neurons sharpen during attentive behavior (Lee and Middlebrooks, 2011). Specifically, the authors observed that A1 neurons exhibited broad spatial tuning when the animal was idle but sharpened their spatial tuning when the animal performed a spatial localization task. We modeled this effect using the AIM network.

In this simulation, the AIM network consisted of an array of spatial channels tuned to locations between −90° and 90° azimuth. We then probed the network with broadband noise from various spatial locations. We found that when I2 neurons in all spatial channels were on, the cortical neuron in the AIM network exhibited broad tuning similar to the idle condition in the experiment (Fig 2A). Release of acetylcholine (ACh) during behavioral task performance can suppress intracortical excitatory connections and strengthen thalamocortical connections (Hasselmo, 1995; Gil et al., 1997; Hsieh et al., 2000). When we simulated these effects in the AIM network, the spatial tuning of the cortical neuron sharpened, resembling the tuning in the behaving condition in the Lee and Middlebrooks study (Figure 2B). In that study, however, animals were not required to attend to a specific location during the task. We simulated selective attention to a specific location by inactivating an I2 neuron in a specific channel, e.g., 30° (Fig 2C, left). We found that, in this case, the spatial tuning of the cortical neuron also sharpened (Figure 2C, right). In the AIM network, this effect occurs because of two key mechanisms: the disinhibition of the attended channel by the I2 neuron, which then drives powerful inhibition of competing channels by the I neuron. Thus, selective attention activates focal disinhibition at the attended location and suppression at other locations in the network.

**Figure 2:**
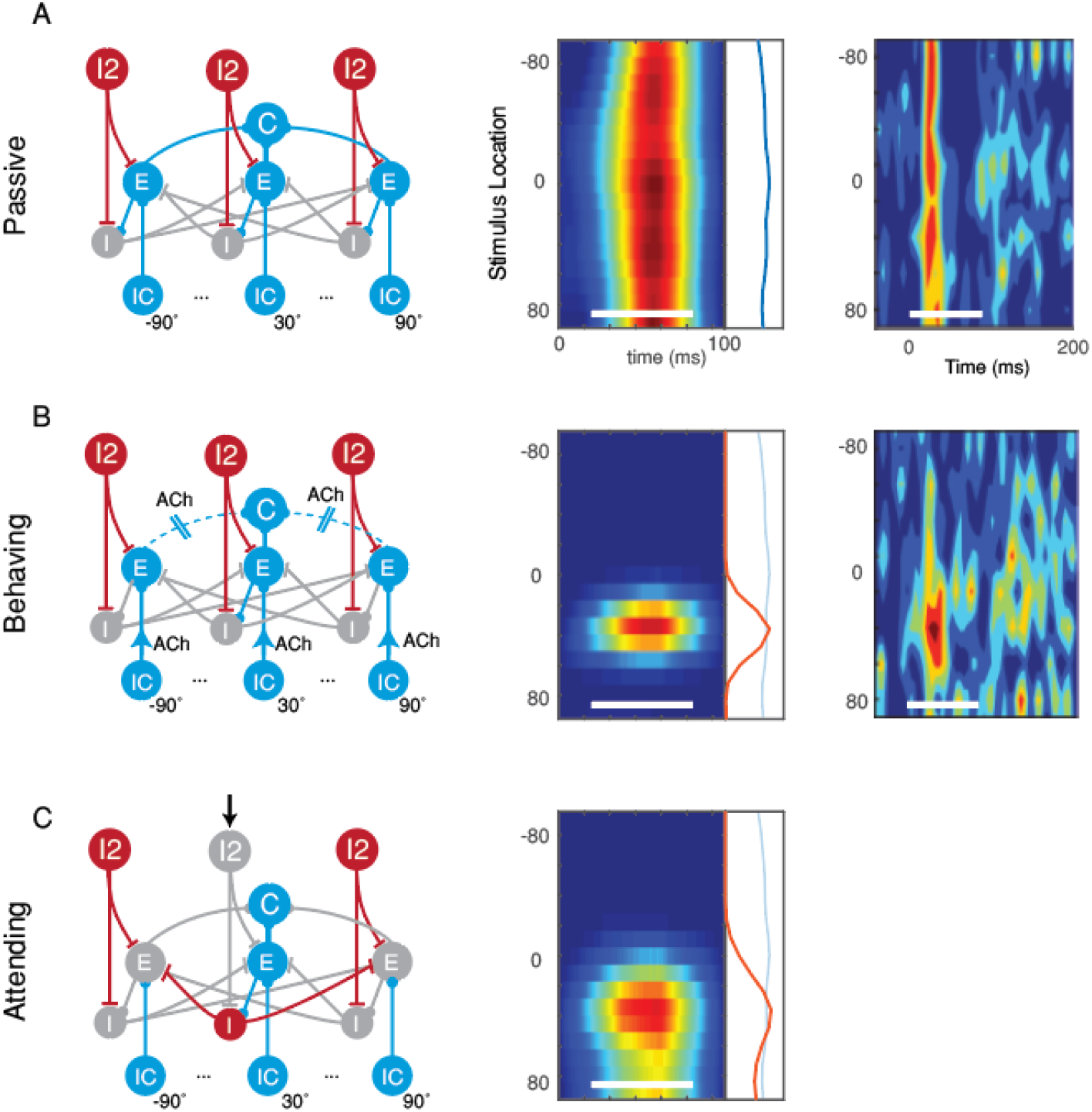
Attentional sharpening of spatial tuning. First column shows the spatial AIM network in (A) passive condition. (B) Behaving condition, where the network is modulated with the cholinergic system. (C) Attending condition, here selective attention is simulated by controlling the state of specific I2 neurons. Second column shows the spatial tuning, i.e., peristimulus time histogram expanded vertically to show the spatial dimension, calculated from the network output. Firing rate is shown (red is higher) as a function of stimulus location over time, in response to a broad band noise. White bar indicates the duration of the noise stimulus. Third column shows experimentally recorded spatial tuning in the cat A1. Figures from Lee and Middlebrooks reproduced here with the permission of Nature Neuroscience.

**Figure 3:**
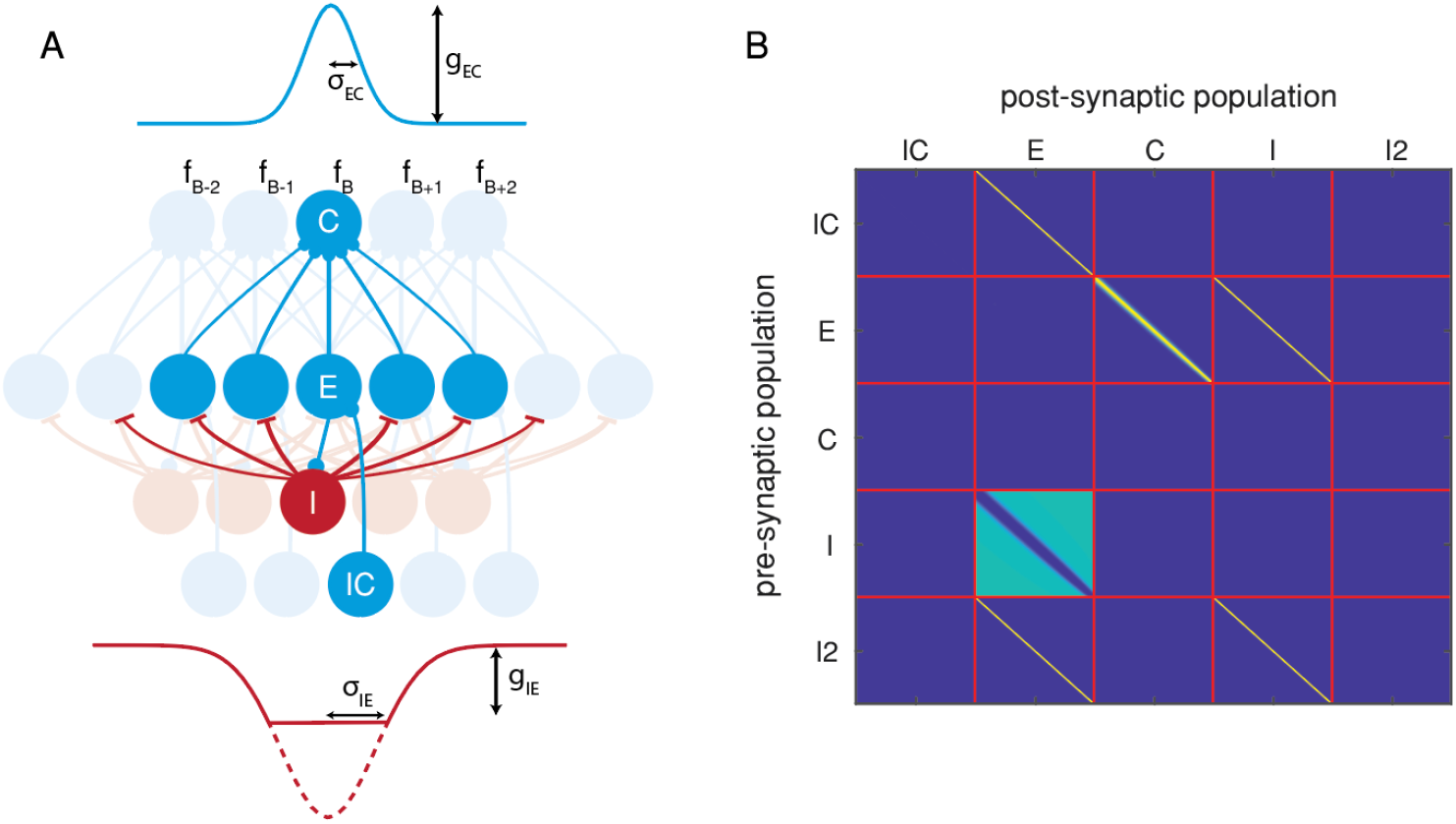
AIM Network Diagram for the spectral network. A) convergence of connectivity across neuron types. E neurons converge to C neurons locally, centered around a neuron at the best frequency, f_B_. The connectivity strength decays across adjacent channels, and is modeled by a Gaussian function. The convergence width is determined σ_EC_, and the connectivity strength is determined by g_EC_ (see table 2). The spread of inhibitory connection from I neurons to E neurons is modeled by a thresholded, inverted Gaussian function. The connectivity within Zσ_ļE_ of this Gaussian function is zero. The inhibition gradually rises to a value determined by g_IE_. B) The network connectivity matrix describes all connections within the network for these simulations.

### Attentional Changes in Spectral Tuning

Rapid changes in receptive fields during task performance, thought to arise from attentional mechanisms, have also been observed in the frequency domain as in the experiments by Fritz et al., (2003) and Atiani et al., (2009). Here, we show that the attentional mechanisms in the AIM network can also account for these experimental observations.

We first constrained the AIM network to incorporate experimentally observed features of network connectivity in the frequency domain (Fig 4). A key difference from the spatial domain is that connectivity across frequency channels is localized to nearby frequencies, reflecting the tonotopic organization, unlike the broad connectivity in the spatial domain (Figs 1,2).

**Figure 4:**
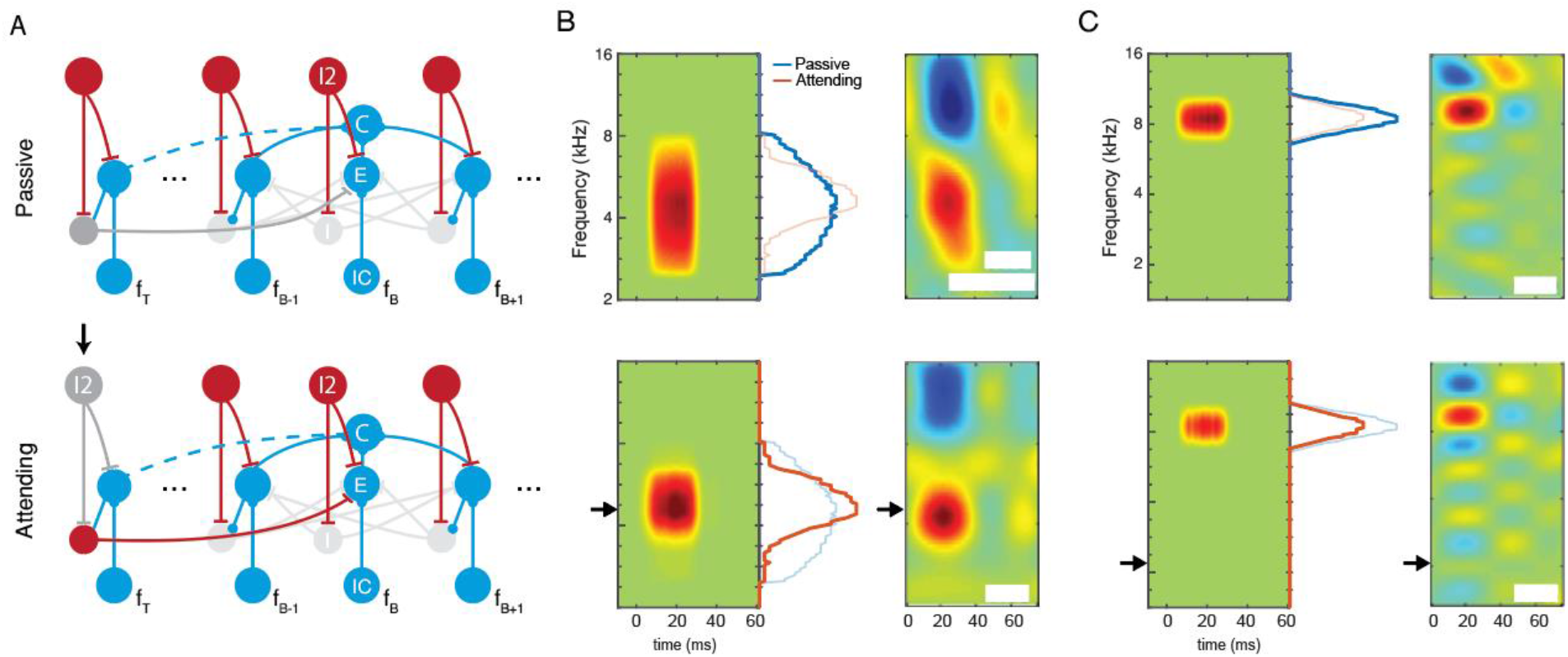
Simulating effects in Atiani et al. A) The AIM network in passive and attending state. Black arrow marks the effect of attention. B) The results of the AIM network (left panels) versus the results described in Atiani et al. (right panels), when f_τ_ (black arrow) is near f_B_. Marginals show the total spiking activity across the duration of the simulation. C) The results of the AIM network versus the results shown by Atiani et al., when f_τ_ is far from f_B_. Figures from Atiani et al. are reproduced here with the permission of Neuron. White patches hide irrelevant text.

We then simulated the spectral tuning of neurons in the network in the passive vs. attentive states. Here, we probed the network with pure-tone stimuli in the frequency ranges shown in Figure 4. In the attentive state, the network attended to a specific target frequency, distinct from the best frequency of the neuron, as in the experiments by Atiani et al.

In the attentive state, the I2 neuron in the attended channel (i.e., the target frequency channel, *f_T_,* Figure 4) is suppressed during attention, disinhibiting the E and I neurons in that channel (Figure 4A). In this case, we found that when the target frequency was close to the best frequency of the neuron, attention produced an increase as well as a sharpening of the response near the best frequency, as observed experimentally (Figure 4B). In the AIM network, this occurs because when the target frequency is close to the best frequency, excitation from the disinhibited E neuron in the target channel increases the peak response of the cortical neuron, and inhibition via the I neuron in the target channel sharpens the shape of response. In contrast, when the target frequency is far from the best frequency of the neuron, the effect due to inhibition dominates, producing a net suppressive effect on the response, which is also observed experimentally (Figure 4C). Thus, the AIM network qualitatively explains salient features of the experimental observations by Atiani et al.

In addition to sharpening, strengthening, and weakening of STRF hotspots, the experimental results of Fritz et al. showed that a secondary hotspot in the STRF may arise when the animal is in the attentive state. (Fig 5).

**Figure 5.**
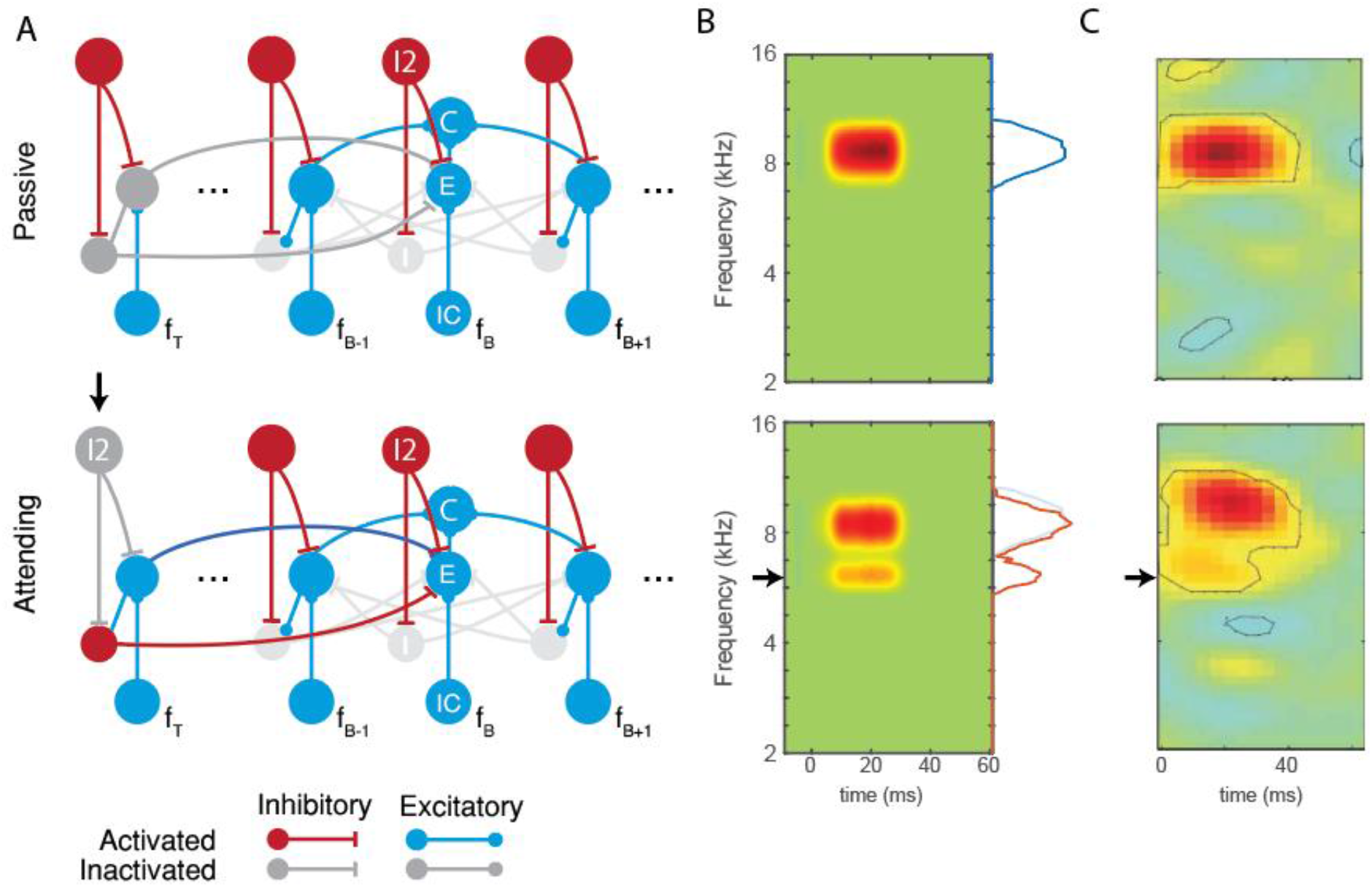
Simulating effects in Fritz et al., A) The proposed mechanism underlying the changes in A1 behavior. Frequency channels adjacent to the best frequency f_B_ as well as the target frequency f_T_ are shown. Black arrow indicates the target of selective attention. The dark blue connection highlights the additional intracortical connection unique to this simulation. B) Simulated A1 neuron STRFs in the two states of attention. The model qualitatively reflects the changes observed in physiological recordings. C) Physiological recording of a STRF of an A1 neuron when the animal is passively listening (top) and when the animal is attending to a target tone, marked by the black arrow (bottom). Figures from Fritz et al. are reproduced here with the permission of Nature Neuroscience.

We hypothesized that this is a result of the strengthening of an intracortical connection between the target frequency and the best frequency in the attentive state (see *Discussion* for possible mechanisms). To test this hypothesis, we added an additional E-E connection between the *f_τ_* and *f_B_*, and found that in the passive state, the neuron responded to frequencies near its best frequency, showing a single hotspot. On the other hand, in the attentive state, the same neuron also responded to the target frequency, as seen by the emergence of a new excitatory region at the target frequency in Fig 5B. This effect occurs due to the strengthening of synaptic connection between the target frequency and the best frequency in the attentive state. There is also a suppressive effect on the response to the best frequency, as seen in the slight reduction in amplitude of the tuning curve at best frequency. This effect occurs due to inhibition driven by the I neuron in the target frequency channel. Both of these effects were observed in experimentally measured receptive fields (Fritz et al., 2003). Thus, the AIM network qualitatively explained salient features of experimentally observed changes by Fritz et al.

### Functional Implications

We hypothesized that the attentional mechanisms in the AIM network play an important functional role in processing complex auditory scenes. Two highly effective mechanisms for sound segregation are spatial hearing and frequency selectivity. We first considered functional implications in the spatial domain.

When entering a cocktail party, we might want to monitor the entire scene, focus our attention on a conversation partner, or switch attention between conversation partners. How does top-down attentional control modulate bottom-up mechanisms to enable this flexible behavior? To illustrate the behavior of the AIM network in these different modes, we simulated several scenarios. In these simulations, we used three spatial channels corresponding 0°, 90°, and −90° azimuths, presenting the network with different tokens of speech stimuli at these locations, either sequentially or simultaneously (see Methods).

To demonstrate passive listening (the “monitor” mode), we set all I2 neurons active, thereby silencing all I neurons (Figure 6A). In this case, when the speech tokens were presented sequentially to the network, the network output resembles each individual speaker (Figure 6A). When the speech tokens were presented simultaneously, the network output resembles their mixture. Thus, in this mode, the network broadly monitors the acoustic scene across different spatial locations.

**Figure 6:**
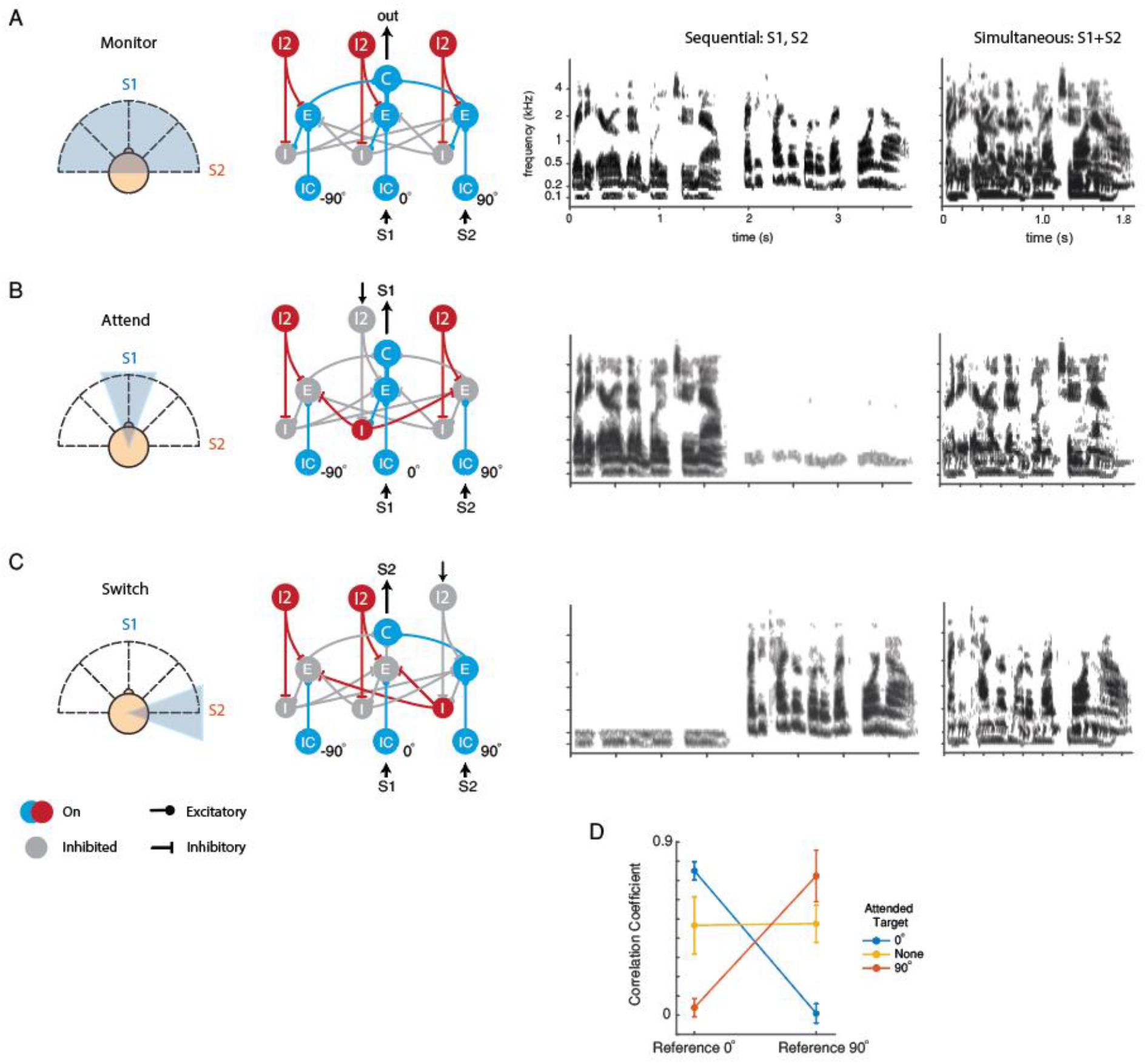
Functional implications of the AIM network: Spatial tuning of the network is dictated by the state of TD neurons. (A) The network monitors the entire azimuthal plane when all TD neurons are active. (B) The network attends to a specific direction if the corresponding TD neuron is off. (C) The network attends to a different location if a different TD neuron becomes inactive. Column 3 shows the result of simulations when speakers are presented sequentially to the network, in spike rasters. Column 4 shows the result of simulations when speakers are presented simultaneously to the network. (D) Cross correlation measures between the simultaneous simulation results and single speaker spike rasters in the passive condition show that the network can “attend” to particular location in space. Error bars show standard deviation (n = 20). X-axis is the reference speaker, and each line color denotes the attended location.

To simulate selective attention to a particular speech token, we first inactivated the I2 neuron in the 0° spatial channel, thereby activating lateral inhibition via the disinhibition of I neurons in that channel (Figure 6B). In this case, the network output resembled the 0° speaker output, regardless whether the speakers were presented sequentially or simultaneously. Finally, to simulate switching attention to a different location (90°), we turned off the I2 neuron at 90°, activating lateral inhibition via the disinhibition of I neuron at that location (Fig 6C). In this case, the network output was more similar to the output for the speaker at 90°. In summary, when an I2 neuron in a specific spatial channel is inactivated, it disinhibits the I neuron at that location, causing the network to selectively attend to that spatial location.

We next investigated the functional role of the AIM network in the frequency domain using the same network shown in Figure 3. In this simulation, two competing speech tokens, a male and female speaker, originate from the same spatial location. In this case, differences in spatial cues cannot be exploited for segregation, but differences in spectral features of the two speakers, e.g., the fundamental frequency (F0), are available. We simulated the network in three conditions: passive, attending to male F0, or attending to female F0. Attention was simulated (as in Figure 2) by inactivating the I2 neuron which disinhibits the I neuron at the attended frequency channel. We found that in the attending modes, network activity showed a sharp peak around the F0 of the attended speaker, and a suppression of activity around the F0 of the competing speaker (Fig 7). Such enhancement of an attended feature accompanied by suppression of competing features may contribute to speech segregation in settings where spatial cues are unavailable (Mesgarani and Chang, 2012).

**Figure 7.**
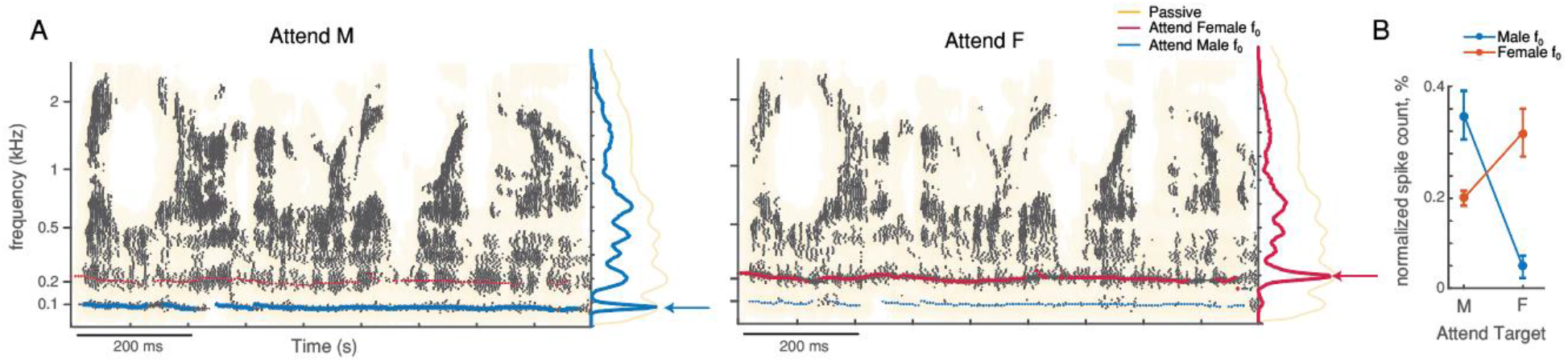
Functional role of the AIM network in the frequency domain. (A) AIM network outputs (raster plots) when attending to the male talker’s f_0_ (left) or female talker’s f_0_ (right). Yellow shaded background shows the spike raster of the network output in the passive condition. Blue and red lines mark the estimate f_0_ for each speaker. Blue and red lines in the marginal show the normalized total spike counts per frequency channel. Yellow line in the marginal show total spike counts of the AIM network in the passive condition. (B). Change in normalized spike count (%) in the male or female f_0_ channel when attending to the male or female target. Error bars show standard deviation (n = 20).

## Discussion

### Attention and Flexible Cortical Processing

The capacity for generating flexible behaviors in a context-dependent manner is central to most complex cognitive tasks. How cortical circuits achieve such flexible computations is a central area of investigation in both theoretical and experimental neuroscience. Cortical mechanisms underlying flexible behaviors should be distinguished from long term synaptic plasticity, implicated in developmental and perceptual learning. Specifically, flexible behaviors require relatively rapid, reversible changes in system outputs in a context-sensitive manner. Recent theoretical studies have begun to propose model networks capable of producing flexible behaviors, e.g., gating mechanisms for flexible routing of information (Vogels and Abbott, 2009; Yang et al., 2016; Wang and Yang, 2018). Experimental studies have begun to reveal cortical mechanisms underlying flexible gating of information and attentional control^4-6^.

Changes in attentional states could be realized via neuromodulatory systems which exert powerful control over the global state of cortical networks, e.g., asleep, quiet arousal, and attending (Lee and Dan, 2012). Two key modulatory projections to cortex involve norepinephrine (NE) and acetylcholine (ACh), which have been implicated in arousal and attention. In our network, we assumed that when the network is in the “monitor” mode, the I2 layer is on. Such a global state may correspond to arousal and be NE-dependent. Top-down suppression of I2 neurons in our model could correspond to an attentive state and be ACh-dependent. Although cholinergic mechanisms are necessary for modulating the state of the network, these mechanisms alone are not sufficient to achieve selective attention.

The flexibility of the AIM network is generated by top-down inputs, which control the state of the network by dictating the on/off state of specific I2 neurons. In the spatial case, when I2 neurons in all channels are on, the network integrates information from all input channels, producing broad spatial tuning and allowing the network to monitor the entire scene. This is important from a functional standpoint. A network that is always selective for a single channel may fail to detect important events in the scene at other locations. The different states of the I2 layer (e.g., all on vs. one off), allows the opportunity for *exploration* (by detecting events across a broad range of locations), as well *exploitation* (by selectively listening to a particular channel), in a dynamic manner. This is especially interesting in the context of findings that the “spotlight of attention” fluctuates in a rhythmic manner (Helfrich et al., 2018). In our model, such fluctuations in the strength of the top-down input would cause the network to alternate between periods favoring broad detection of sounds across the entire acoustic scene and fine discrimination at a single spatial location (Guo et al., 2017). Such periods may allow salient sounds in the background to capture the spotlight of attention, as demonstrated in a recent study in humans (Huang and Elhilali, 2020). Moreover, changes in the location of the top-down input would promote switching attention to a different location (Fiebelkorn and Kastner, 2019)

### Cortical Inhibitory Neurons and Function

Inhibitory neurons play key roles in the AIM network. There are several types of inhibitory neurons in cortex (Tremblay et al., 2016). The majority of inhibitory neurons can be placed into three categories: those that express parvalbumin (PV), somatostatin (SOM), or vasointestinal peptide (VIP). It is worth noting that most currently available information on specific classes of interneurons come from studies in rodents in a variety of cortical areas, whereas key experimental observations modeled in this study were obtained in other species. Thus, it is difficult to directly map the functional groups of neurons, e.g., I2 and I, in the model to identified interneuron types e.g., PV, SOM and VIP neurons. Nevertheless, we suggest some hypotheses on a possible correspondence based on recent experiments in rodents, to motivate future experimental work.

The top-down input in the model could correspond to inputs from VIP neurons. VIP mediated inhibition is engaged under specific behavioral conditions, including attention (Pi et al., 2013; Zhang et al., 2014). It has been proposed that VIP cells “open holes in the blanket of inhibition” (Karnani et al., 2016), generating the “spotlight of attention” (Zhang et al., 2014). The model is consistent with this intuition, with the top-down input being critical for selecting a particular target and switching to a different target. VIP input is often thought to favor excitation, due to the disinhibition of excitatory neurons (Pfeffer, 2014). In the model, top-down inhibition of an I2 neuron in a specific channel activates powerful inhibition via I neurons that suppress competing channels, leading to the selection of the target. Thus, the model also explains powerful suppressive effects of selective attention, which have been observed in auditory cortex (Otazu et al., 2009). The model predicts that silencing top-down input to a specific channel, via optogenetics or other methods, should block the effects of selective attention.

VIP neurons inhibit SOM neurons, which in turn inhibit excitatory neurons (Pfeffer et al., 2013). This motif suggests that the I2 neurons in our model may correspond to SOM neurons, specifically Martinotti cells, which are strongly targeted by VIP neurons (Muñoz et al., 2017).

The I neurons mediate powerful and sustained inhibition of competing channels in the model. A key distinction between this type of inhibition and “classical” lateral inhibition observed at multiple stages of sensory processing starting at the periphery is noteworthy. Classical lateral inhibition is activated by bottom-up stimulus driven mechanisms, whereas the inhibition in our model is driven by top-down attentional mechanisms. To distinguish these two cases, we refer to the inhibition mediated by I neurons in the model as “top-down lateral inhibition”. Top-down lateral inhibition can be activated by direct disinhibition of I neurons, disinhibition of E neurons which drives feedback lateral inhibition via I neurons (Supplementary Fig), or a combination of the two.

In principle, top-down lateral inhibition could be mediated by any interneuron type. Although PV neurons are a possible candidate, the long-lasting inhibition required to suppress competing channels in our model should be distinguished from the fast and transient dynamics of inhibition typically associated with PV neurons (Tremblay et al., 2016). SOM neurons can also mediate feedback lateral inhibition to generate a “winner-take-all” circuit and suppress competing channels (Silberberg and Markram, 2007), or modulate bottom-up inputs in specific layers (Muñoz et al., 2017; Naka et al., 2019). Developing behavioral paradigms for investigating attention in rodents combined with optogenetic manipulations, and/or developing methods for selectively manipulating different interneuron types in other species, are promising future directions for identifying specific cell-types involved in mediating top-down lateral inhibition.

### Space vs. Frequency

We related the effects of attention in the AIM network to key experimental observations on changes in cortical spatial and frequency tuning in animals engaged in a behavioral task vs. passive animals (Fritz et al., 2003; Lee and Middlebrooks, 2011). Similar changes have also been reported in the primary auditory cortex of humans (van der Heijden et al., 2018). We assume that attentional mechanisms are a key factor in driving such changes (Fritz et al., 2007a, 2007b; Lee and Middlebrooks, 2011). The effects of attention appear very different in the spatial and frequency domains. In the spatial domain, broad tuning sharpens during task performance; whereas in the frequency domain, narrow tuning can be enhanced or suppressed depending on the target frequency and other parameters such as SNR. Our results suggest these apparent differences in the spatial vs. the frequency domain do not require differences in the underlying attentional mechanisms. A key difference between the spatial and frequency domains in our model is the convergence from the E neurons to the C neuron, which is broad in the spatial domain and narrow in the frequency domain. Previous studies in A1 have found a tonotopic organization, but no topographic organization for spatial tuning (Panniello et al., 2018). Local synaptic connections in a patch of cortex may therefore result in convergence from neurons with similar tuning in frequency but a broad range of spatial tuning, consistent with our model. Thus, our results reveal that the same cortical mechanisms underlying attention can produce diverse effects on stimulus tuning, due to differences in the cortical organization of a stimulus feature, e.g., space or frequency.

The key mechanisms underlying selective attention in the AIM network are the top-down inhibition of I2 neurons which disinhibits the attended channel and suppresses competing channels via top-down lateral inhibition, mediated by I neurons. Although these mechanisms explain diverse experimental observations, alternative/additional mechanisms may be involved in some aspects of the experiments modeled here. For the sharpening of spatial tuning, we found that global cholinergic effects on cortical circuits, i.e., suppression of intracortical connections and enhancement of thalamocortical connections, could also produce a sharpening in spatial tuning (Figure 2). In the study by Lee and Middlebrooks (2011), animals were not required to attend to a fixed location when performing the localization task. Thus, activation of global cholinergic mechanisms may be more consistent with that experimental design. Our model predicts that experiments where animals are required to selectively attend to a specific location should also produce a sharpening of spatial tuning in A1. For the experiments by Fritz et al. (2003), we found that strengthening of the intra-cortical synaptic connection between the target frequency and the best frequency could explain the emergence of new excitatory regions at the target frequency. An interesting possible mechanism for transient, reversible strengthening of intracortical synapses is synapse specific gating (Yang et al., 2016; Wang and Yang, 2018), which may then promote long-term strengthening via classic Hebbian plasticity.

The model has several simplifications and limitations that motivate future directions of work. For example, in the frequency network, we used pure tones to characterize the responses of neurons in the network and relate them to experimentally observed STRFs obtained with ripple noise stimuli. Although this approach captured salient attentional effects observed experimentally, future studies should probe non-linear components using complex stimuli, e.g., ripples and natural sounds, and fully characterize the structure of STRFs including inhibitory subregions. For simplicity, we considered spatial and spectral networks separately. Future models should unify these two dimensions. Here, we did not explicitly model how spatially tuned inputs to the AIM network arise, an aspect that is likely to be species dependent. In the AIM network, spatial tuning is inherited from tuning for acoustic cues in pre-cortical areas (Chou et al., 2019), perhaps the simplest scenario consistent with experimental observations (Knudsen and Konishi, 1978; Yin and Chan, 1990; Köppl and Carr, 2008). Additional mechanisms, e.g., forward suppression, may further sharpen or generate spatial tuning in cortex (Zhou and Wang, 2014; Yao et al., 2015). In rodents, spatially tuned responses covering a range of azimuths have been observed in cortical areas (Higgins et al., 2010), and may emerge from excitatory-inhibitory interactions in the underlying network (Kyweriga et al., 2014). From a functional standpoint, it is interesting to note that sharp tuning is not necessary for the monitor mode, but only for the selective mode of the AIM network. In some species, the sharpness of tuning may emerge in the attentive state based on state-dependent mechanisms, and/or inputs from other brain areas, e.g., the superior colliculus, which shows a map of auditory space (Middlebrooks and Knudsen, 1984; King and Hutchings, 1987) and can modulate responses in A1 via the pulvinar (Chou et al., 2020). These outstanding issues will require further experimental work, especially in attentive animals, as well as the development of species-specific models. Finally, we did not model auditory streaming, an important class of phenomena in auditory attention. We conjecture that modeling streaming at the cortical circuit level will require modeling rich temporal aspects of neuronal and network dynamics, e.g., adaptation, synaptic facilitation and depression, oscillations, synchrony and coherence. Future extensions of the AIM network should incorporate these aspects to link mechanisms of neuronal and network dynamics to attentional dynamics.

### Models of Attention

Previous studies have modeled the effects of attention on auditory cortical receptive fields using mathematical and computational principles, e.g., temporal coherence (Krishnan et al., 2014; Kaya and Elhilali, 2017). In contrast, the AIM network is a cortical circuit level model underlying attentional effects. One recent study modeled different STRFs in the attending vs passive state of the ferret A1 with a two-layer spiking network (Chambers et al., 2019). The focus of that study was to produce detailed fits of STRFs in attending and passive animals. In contrast, the focus of this study was to propose general cortical circuit mechanisms, e.g., top-down disinhibition, underlying the effects of attention on both spatial and spectral tuning. Another previous study modeled global cholinergic mechanisms underlying changes in STRF (Soto et al., 2006), similar to the effects modeled in Figure 2B. However, that study did not include the selective top-down disinhibitory mechanism, which was unknown at that time, and is a key mechanism in the AIM network.

Original models of attention in vision were also developed based on computational principles, e.g., biased competition or normalization (Desimone and Duncan, 1995; Reynolds and Heeger, 2009). At the time, available information on cortical circuits to guide and constrain circuit-based models were limited. Subsequently, cortical circuit-based models of visual attention have been proposed (Ardid et al., 2007). With the rapidly emerging knowledge of specific cell types and circuitry in auditory cortex, and the availability of powerful optogenetic tools for cell type-specific perturbations, the AIM network may help guide the design of new experiments to unravel cortical circuits that underlie auditory attention.

## Methods

Simulations and models were implemented in Matlab (Natick, MA). Code for the AIM network is available upon request.

## Stimuli

Three sets of auditory stimuli were used, depending on the particular simulation. White Gaussian noise was used as the stimulus in spatial tuning simulations, pure tones with frequencies approximately equal to the center frequencies of the gammatone filterbank (see Subcortical Processing) were used in spectral tuning simulations, and speech tokens from the Coordinated Response Measure (CRM) corpus were used in the functional demonstration simulations (Bolia et al., 2000). In spatial simulations where stimuli were placed along the azimuth, directionality is imparted on the stimuli by convolving them with the head-related transfer functions (HRTFs) of the Knowles Electronics Mannikin for Acoustic Research (KEMAR) (Burkhard and Sachs, 1975; Algazi et al., 2001).

## Subcortical Processing

Stimuli for each simulation were first processed and encoded with models of the auditory periphery and midbrain, then presented to the network. The auditory periphery was modeled by a gammatone filterbank, implemented using the Auditory Toolbox (Slaney, 1998). It was used to separate the sentence mixture into 64 narrowband frequencies, with center frequencies ranging from 200 to 8000 Hz, uniformly spaced on the equivalent rectangular bandwidth scale.

We used a previously published model of the midbrain to perform spatial segregation of spatialized stimulus mixtures, as well as to encode the stimuli. If a simulation did not use spatial stimuli as the input, the stimuli were treated as dichotic. For details pertaining to the midbrain model, see Fischer et al., (2009) and Chou et al., (2019). Briefly, the midbrain model computed binaural features (i.e., interaural timing and level differences) in each time-frequency tile (i.e., narrowband and short time window). Model neurons encoded the stimulus at specific timefrequency tiles if the binaural features of the stimuli matched the “preferred” binaural features of the model neuron, thereby performing spatial segregation. The preferred binaural feature of each model neuron is specific to the frequency and spatial channel each neuron belonged to. There were 64 frequency channels in the midbrain model, corresponding to each channel of the gammatone filter. The number of spatial channels in the midbrain model depended on each specific simulation. The input neuron in a spatial channel is spatially tuned to the azimuth corresponding to that channel, consistent with spatial tuning of acoustic cues observed in sub-cortical areas (Knudsen and Konishi, 1978; Yin and Chan, 1990; Köppl and Carr, 2008). The spiking responses of these model neurons were used as the input to the AIM network.

## Attentional Inhibitory Modulation (AIM) Network

The AIM network was implemented using the DynaSim package (Sherfey et al., 2018), and its structure is illustrated in Figure 1. For simplification purposes, only one frequency channel and three spatial channels are shown. A “spatial channel” refers to the sub-network of neurons that are responsible for processing inputs from a specific spatial location (blue shading, Figure 1). The number of spatial and frequency channels in the network, and their connectivities, depended on the specific simulation being explored.

Five neural populations were created within the network: excitatory input (IC), excitatory (E), inhibitory (I), output cortical (C), and a second inhibitory (I2) population. IC neurons represent the bottom-up inputs to the network from the subcortical model. I2 neurons represent attentional top-down control. With the exception of the C neurons, a number of neurons were created within each population, corresponding to each of the spatial or frequency channels needed in a simulation. All five neural populations are implemented as leaky integrate-and-fire neurons whose dynamics are defined by Dayan and Abbott (2001):

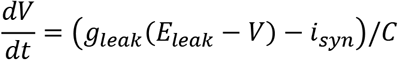

Where *V* is the membrane potential, *i_tonic_* is tonic firing current, *i_syn_* is the synaptic input current, *C* is membrane capacitance, *g_leak_* is the membrane conductance, *E_leak_* is the equilibrium potential, *V_thresh_* is the action potential threshold, *V_spike_* is the action potential voltage, and *V_reset_* is the reset voltage. Values for these parameters are listed in Table 1. The dynamics of the synaptic input current is defined by a double exponential:

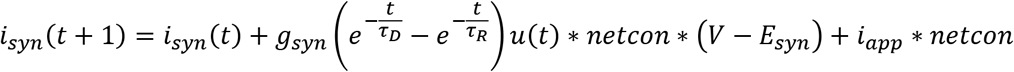

**Table 1:** Default parameters of cellular dynamics

| Parameter | Value |
| --- | --- |
| $C$ (nF) | 1 |
| $g_{leak}$ ($\mu S$) | 1/10 |
| $E_{leak}$ (mV) | -70 |
| $V_{thresh}$ (mV) | -55 |
| $V_{spike}$ (mV) | 50 |
| $V_{reset}$ (mV) | -75 |
| $i_{tonic}$ ($\mu A$) | 0 |
| Noise | 0 |

Where *t* is time, *g_syn_* is the synaptic conductance, *τ_D_* and *τ_R_* are the decay and rise time constants, respectively, *u(t)* is the unit step function, *E_syn_* is the reversal potential, *i_e_* is the externally applied current, and *netcon* refers to a binary matrix that defines the connections between neurons. Inhibitory synapses have the following parameters: *τ_R_* = 1*ms*, *τ_D_* = 10*ms*, *E_syn_* = −80*mv*. Excitatory synapses have the following parameters: *τ_R_* = 0.4*ms*, *τ_D_* = 2*ms*, *E_syn_* = 0*mv*. The values for *g_syn_* and *i_app_* are simulation- and connection-dependent, and are listed in Table 2. The network connections are illustrated in Figures 1 and 3.

**Table 2:** Simulation-specific parameters. *g_syn_* have units of *μS* and *i_app_* have units of *nA.* Parameters of local convergence *g_IE_*, *g_EC_ σ_IE_*, and *σ_EC_* are also shown (Figure 3).

| Connection<br>or Neuron | Param | Lee | Fritz | Atiani a | Atiani b | Spatial<br>Function | Freq<br>Function |
| --- | --- | --- | --- | --- | --- | --- | --- |
| $IC \rightarrow E$ | $g_{SYN}$ | 2.5 | 4 | 4 | 3 | 2 | 4 |
| $E \rightarrow C$ | $g_{SYN}$ | 2 | 1.25 | 1.25 | 3 | 2 | 1.5 |
| $E \rightarrow I$ | $g_{SYN}$ | 2.5 | 3 | 3 | 3 | 2 | 3 |
| $E \rightarrow E$ (Attend) | $g_{SYN}$ | - | 5 | - | - | - | - |
| $E \rightarrow E$ (Passive) | $g_{SYN}$ | - | 3 | - | - | - | - |
| $I2 \rightarrow I$ | $g_{SYN}$ | 4 | 3 | 3 | 3 | 2.25 | 3 |
| $I2 \rightarrow E$ | $g_{SYN}$ | 2.8 | 4 | 4 | 3 | 1.25 | 3 |
| $I \rightarrow E$ | $g_{SYN}$ | 3 | 4 | 4 | 4 | 3 | 3.5 |
| $I$ | $i_{app}$ | 3 | - | - | - | 4 | - |
| $E$ | $i_{app}$ | 1 | 3 | 3 | 3 | 0 | 3 |
| $I2$ | $i_{app}$ | 8 | 8 | 8 | 8 | 3.5 | 8 |
| $g_{IE}$ | - | - | 0.5 | 0.5 | 0.175 | - | 0.175 |
| $g_{EC}$ | - | - | 1 | 1 | 1 | - | 1 |
| $\sigma_{IE}$ (kHz) | - | - | 0.65 | 1.7 | 2 | - | 0.01 |
| $\sigma_{EC}$ (kHz) | - | - | 0.32 | 1.15 | 0.35 | - | 0.05 |

The default *g_syn_* were chosen such that if I neurons were off, then the inputs would be relayed and combined at the C neuron with a similar firing rate, and if I neurons were on, then E neurons would be completely silenced.

## Simulation-Specific Model Configurations

### Lee & Middlebrooks simulations

In this spatial tuning simulation, 80 ms of white gaussian noise was placed between −80° to 80° azimuth, in 10° increments. The spatialized stimuli were then processed and encoded with the subcortical model. The midbrain model in this simulation consisted of 19 spatial channels from −90° to 90° azimuth, in 10° increments, and 64 frequency channels. To reduce the computational demand of simulating the AIM network, a new set of spike trains, generated using a Poisson model based on the overall firing rate across all frequency channels, were computed for each spatial channel. This operation essentially collapses the neural response over the frequency dimension. Therefore, the AIM network for this simulation consisted of 19 spatial channels and one single frequency channel, where each spatial channel processed the set of spike trains that represent the average activity across all frequencies. Network connectivities between spatial channels are as shown in Figure 2. Spatial tuning curves were then calculated based on the response of the C neuron of the AIM network.

The effects of neuromodulators were simulated by applying a gain on the network connections. During the behavior state, off-target E-C connectivities were applied a gain of 0 to simulate the effects of muscarinic receptors, and off-target IC-E connectivities were applied a gain of 2.5 to simulate the effects of the nicotinic receptors.

### Atiani et al. and Fritz et al. simulations

Pure tones were presented dichotically to the subcortical model, which consisted of a single spatial channel, corresponding to 0° azimuth, and 64 frequency channels. Spike trains were passed directly to the AIM network, which also consisted of a single spatial channel and 64 frequency channels. Network connectivities between spatial channels are as shown in Figure 3. Spectral temporal receptive fields were calculated based on the response of the best-frequency cortical neuron to each of the pure tone stimulus.

### Functional example – spatial listening

20 pairs of speech tokens, one male and one female, were randomly chosen from the CRM corpus, and placed at 0° and 90° azimuth. For simultaneous presentation, speech tokens were summed prior to being processed by the subcortical model. The subcortical model used 5 spatial channels, tuned from −90° to 90° azimuth in 45° increments, and 64 frequency channels, and spatially segregates the speech tokens. Its output is relayed directly to the AIM network, which also has 5 spatial channels and 64 frequency channels. In this simulation, network connectivities across spatial channels are as shown in Figure 2, and each frequency channel operated independently of each other.

### Functional example – monaural listening

The same 20 pairs of speech tokens as above were summed and presented dichotically to the subcortical model. Here, both the subcortical model and the AIM network has one spatial channel (0° azimuth) and 64 frequency channels. In this simulation, the network connectivities are as shown in Figure 3. The pitch of each speech token was estimated using MATLAB Audio Toolbox’s pitch() function.

## Acknowledgements

The authors thank Larry Abbott and John Middlebrooks for comments on the manuscript. This work was supported by an NSF Award 1835270

## Supplementary Material

**Supplementary figure 1:**
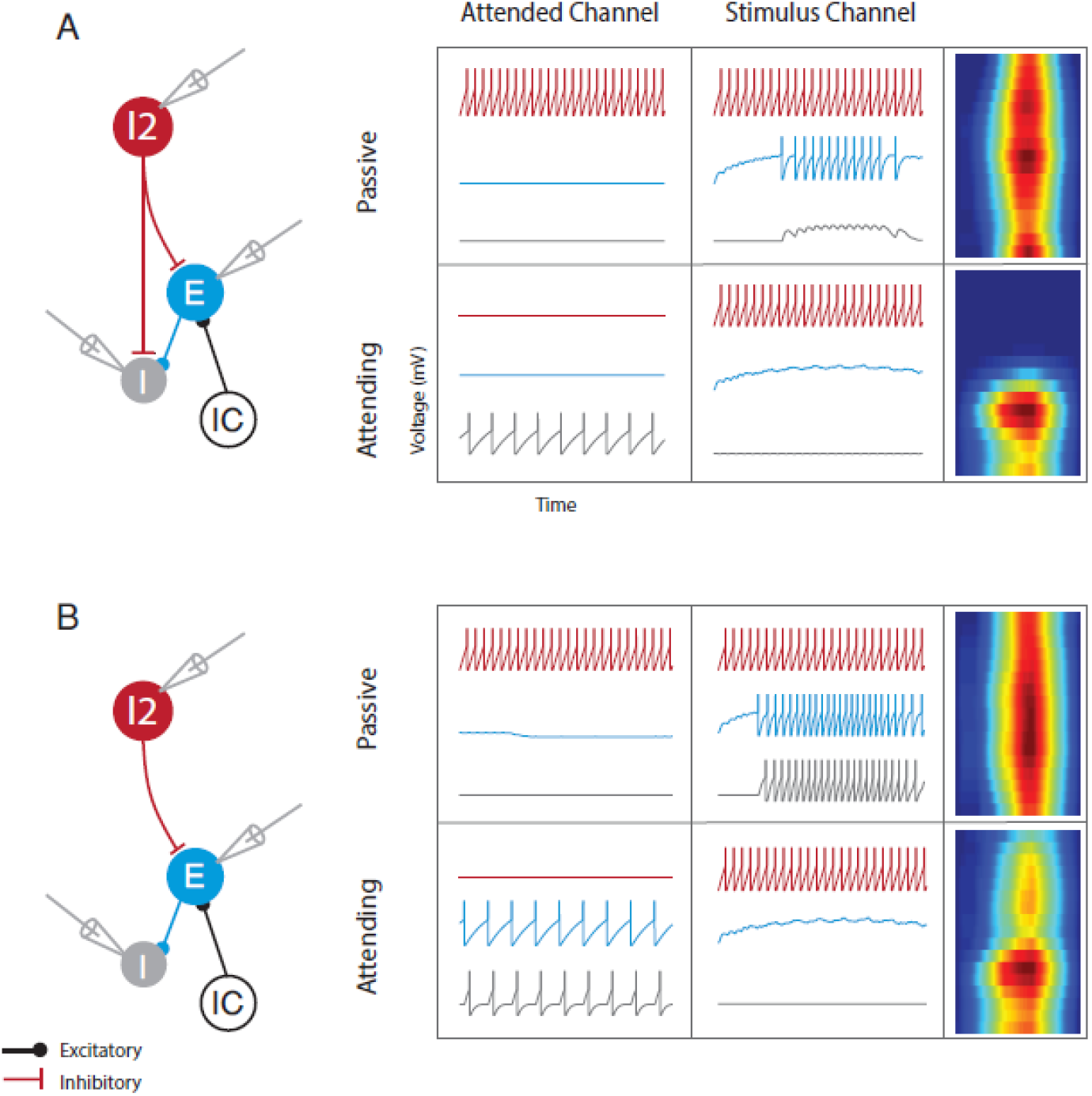
Two possible modes to achieve top-down lateral inhibition, resulting in sharpening of the spatial tuning of the attended channel. A) Direct inhibition from I2 to I neuron, and B) Feed forward inhibition from I2 -> E -> I neuron. Voltage traces of each model neuron under the passive or attending condition is shown on first two columns of the grid. The final column shows spatial tuning of the attended channel in the passive vs attending conditions.

## References

Algazi VR, Duda RO, Thompson DM, Avendano C (2001) The CIPIC HRTF database. In: Proceedings of the 2001 IEEE Workshop on the Applications of Signal Processing to Audio and Acoustics (Cat. No.01TH8575), pp 99–102. IEEE.

Ardid S, Wang XJ, Compte A (2007) An integrated microcircuit model of attentional processing in the neocortex. J Neurosci.

Atiani S, Elhilali M, David S V., Fritz JB, Shamma SA (2009) Task Difficulty and Performance Induce Diverse Adaptive Patterns in Gain and Shape of Primary Auditory Cortical Receptive Fields. Neuron 61:467–480.

Bee MA, Micheyl C (2008) The cocktail party problem: What is it? How can it be solved? And why should animal behaviorists study it? J Comp Psychol 122:235–251.

Bolia RS, Nelson WT, Ericson MA, Simpson BD (2000) A speech corpus for multitalker communications research. J Acoust Soc Am 107:1065–1066.

Bronkhorst AW (2015) The cocktail-party problem revisited: early processing and selection of multi-talker speech. Attention, Perception, Psychophys 77:1465–1487.

Buffalo EA, Fries P, Landman R, Liang H, Desimone R (2010) A backward progression of attentional effects in the ventral stream. Proc Natl Acad Sci U S A.

Burkhard MD, Sachs RM (1975) Anthropometric manikin for acoustic research. J Acoust Soc Am 58:214–222.

Chambers JD, Elgueda D, Fritz JB, Shamma SA, Burkitt AN, Grayden DB (2019) Computational neural modeling of auditory cortical receptive fields. Front Comput Neurosci 13:1–13.

Chou KF, Dong J, Colburn HS, Sen K (2019) A Physiologically Inspired Model for Solving the Cocktail Party Problem. J Assoc Res Otolaryngol 20:579–593.

Chou XL, Fang Q, Yan L, Zhong W, Peng B, Li H, Wei J, Tao HW, Zhang LI (2020) Contextual and cross-modality modulation of auditory cortical processing through pulvinar mediated suppression. Elife 9:1–21.

Dayan P, Abbott LF (2001) Theoretical neuroscience: computational and mathematical modeling of neural systems. Cambridge, MA: MIT Press.

Desimone R, Duncan J (1995) Neural mechanisms of selective visual attention. Annu Rev Neurosci.

Dong J, Colburn HS, Sen K (2016) Cortical Transformation of Spatial Processing for Solving the Cocktail Party Problem: A Computational Model. eNeuro 3:1–11.

Fiebelkorn IC, Kastner S (2019) A Rhythmic Theory of Attention. Trends Cogn Sci.

Fischer BJ, Anderson CH, Peña JL (2009) Multiplicative auditory spatial receptive fields created by a hierarchy of population codes. PLoS One 4:24–26.

Fritz J, Shamma S, Elhilali M, Klein D (2003) Rapid task-related plasticity of spectrotemporal receptive fields in primary auditory cortex. Nat Neurosci 6:1216–1223.

Fritz JB, Elhilali M, David S V., Shamma SA (2007a) Does attention play a role in dynamic receptive field adaptation to changing acoustic salience in A1? Hear Res 229:186–203.

Fritz JB, Elhilali M, David S V, Shamma SA (2007b) Auditory attention—focusing the searchlight on sound. Curr Opin Neurobiol 17:437–455.

Gil Z, Connors BW, Amitai Y (1997) Differential regulation of neocortical synapses by neuromodulators and activity. Neuron.

Guo W, Clause AR, Barth-Maron A, Polley DB (2017) A Corticothalamic Circuit for Dynamic Switching between Feature Detection and Discrimination. Neuron.

Hasselmo ME (1995) Neuromodulation and cortical function: modeling the physiological basis of behavior. Behav Brain Res.

Helfrich RF, Fiebelkorn IC, Szczepanski SM, Lin JJ, Parvizi J, Knight RT, Kastner S (2018) Neural Mechanisms of Sustained Attention Are Rhythmic. Neuron.

Higgins NC, Storace DA, Escabi MA, Read HL (2010) Specialization of Binaural Responses in Ventral Auditory Cortices. J Neurosci 30:14522–14532.

Hsieh CY, Cruikshank SJ, Metherate R (2000) Differential modulation of auditory thalamocortical and intracortical synaptic transmission by cholinergic agonist. Brain Res.

Huang N, Elhilali M (2020) Push-pull competition between bottom-up and top-down auditory attention to natural soundscapes. Elife.

Hubel DH, Henson CO, Rupert A, Galambos R (1959) “Attention” Units in the Auditory Cortex. Science (80-) 129:1279–1280.

Karnani MM, Jackson J, Ayzenshtat I, Sichani XH, Manoocheri K, Kim S, Yuste R (2016) Opening holes in the blanket of inhibition: Localized lateral disinhibition by vip interneurons. J Neurosci 36:3471–3480.

Kaya EM, Elhilali M (2017) Modelling auditory attention. Philos Trans R Soc B Biol Sci 372.

King AJ, Hutchings ME (1987) Spatial response properties of acoustically responsive neurons in the superior colliculus of the ferret: A map of auditory space. J Neurophysiol 57:596–624.

Knudsen E, Konishi M (1978) A neural map of auditory space in the owl. Science (80-) 200:795–797.

Köppl C, Carr CE (2008) Maps of interaural time difference in the chicken’s brainstem nucleus laminaris. Biol Cybern 98:541–559.

Krishnan L, Elhilali M, Shamma S (2014) Segregating Complex Sound Sources through Temporal Coherence Lewicki M, ed. PLoS Comput Biol 10:e1003985.

Kuchibhotla K V., Gill J V., Lindsay GW, Papadoyannis ES, Field RE, Sten TAH, Miller KD, Froemke RC (2017) Parallel processing by cortical inhibition enables context-dependent behavior. Nat Neurosci 20:62–71.

Kyweriga M, Stewart W, Cahill C, Wehr M (2014) Synaptic mechanisms underlying interaural level difference selectivity in rat auditory cortex. J Neurophysiol 112:2561–2571.

Lee C-CC, Middlebrooks JC (2011) Auditory cortex spatial sensitivity sharpens during task performance. Nat Neurosci 14:108–114.

Lee SH, Dan Y (2012) Neuromodulation of Brain States. Neuron 76:209–222.

Letzkus JJ, Wolff SBE, Meyer EMM, Tovote P, Courtin J, Herry C, Lüthi A (2011) A disinhibitory microcircuit for associative fear learning in the auditory cortex. Nature 480:331–335.

Maddox RK, Billimoria CP, Perrone BP, Shinn-Cunningham BG, Sen K (2012) Competing sound sources reveal spatial effects in cortical processing. PLoS Biol 10.

McDermott JH (2009) The cocktail party problem. Curr Biol 19:R1024–R1027.

Mesgarani N, Chang EF (2012) Selective cortical representation of attended speaker in multi-talker speech perception. Nature 485:233–236.

Middlebrooks J, Knudsen E (1984) A neural code for auditory space in the cat’s superior colliculus. J Neurosci 4:2621–2634.

Middlebrooks JC, Bremen P (2013) Spatial Stream Segregation by Auditory Cortical Neurons. J Neurosci 33:10986–11001.

Muñoz W, Tremblay R, Levenstein D, Rudy B (2017) Layer-specific modulation of neocortical dendritic inhibition during active wakefulness. Science (80-).

Naka A, Veit J, Shababo B, Chance RK, Risso D, Stafford D, Snyder B, Egladyous A, Chu D, Sridharan S, Mossing DP, Paninski L, Ngai J, Adesnik H (2019) Complementary networks of cortical somatostatin interneurons enforce layer specific control. Elife.

Otazu GH, Tai LH, Yang Y, Zador AM (2009) Engaging in an auditory task suppresses responses in auditory cortex. Nat Neurosci.

Panniello M, King AJ, Dahmen JC, Walker KMM (2018) Local and global spatial organization of interaural level difference and frequency preferences in auditory cortex. Cereb Cortex 28:350–369.

Pfeffer CK (2014) Inhibitory Neurons: Vip Cells Hit the Brake on Inhibition. Curr Biol 24:R18–R20.

Pfeffer CK, Xue M, He M, Huang ZJ, Scanziani M (2013) Inhibition of inhibition in visual cortex: The logic of connections between molecularly distinct interneurons. Nat Neurosci 16:1068–1076.

Pi HJ, Hangya B, Kvitsiani D, Sanders JI, Huang ZJ, Kepecs A (2013) Cortical interneurons that specialize in disinhibitory control. Nature 503:521–524.

Reynolds JH, Heeger DJ (2009) The Normalization Model of Attention. Neuron 61:168–185.

Sherfey JS, Soplata AE, Ardid S, Roberts EA, Stanley DA, Pittman-Polletta BR, Kopell NJ (2018) DynaSim: A MATLAB Toolbox for Neural Modeling and Simulation. Front Neuroinform 12:10.

Shinn-Cunningham BG (2008) Object-based auditory and visual attention. Trends Cogn Sci.

Silberberg G, Markram H (2007) Disynaptic Inhibition between Neocortical Pyramidal Cells Mediated by Martinotti Cells. Neuron.

Slaney M (1998) Auditory toolbox: A Matlab Toolbox for Auditory Modeling Work. Interval Res Corp Tech Rep 10:1998.

Soto G, Kopell N, Sen K (2006) Network architecture, receptive fields, and neuromodulation: Computational and functional implications of cholinergic modulation in primary auditory cortex. J Neurophysiol 96:2972–2983.

Tremblay R, Lee S, Rudy B (2016) GABAergic Interneurons in the Neocortex: From Cellular Properties to Circuits. Neuron 91:260–292.

van der Heijden K, Rauschecker JP, Formisano E, Valente G, de Gelder B (2018) Active Sound Localization Sharpens Spatial Tuning in Human Primary Auditory Cortex. J Neurosci 38:8574–8587.

Vogels TP, Abbott LF (2009) Gating multiple signals through detailed balance of excitation and inhibition in spiking networks. Nat Neurosci 12:483–491.

Wang XJ, Yang GR (2018) A disinhibitory circuit motif and flexible information routing in the brain. Curr Opin Neurobiol 49:75–83.

Yang GR, Murray JD, Wang XJ (2016) A dendritic disinhibitory circuit mechanism for pathway-specific gating. Nat Commun 7.

Yao JD, Bremen P, Middlebrooks JC (2015) Emergence of Spatial Stream Segregation in the Ascending Auditory Pathway. J Neurosci 35:16199–16212.

Yin TCT, Chan JCK (1990) Interaural time sensitivity in medial superior olive of cat. J Neurophysiol 64:465–488.

Zhang S, Xu M, Kamigaki T, Hoang Do JP, Chang W-C, Jenvay S, Miyamichi K, Luo L, Dan Y (2014) Long-range and local circuits for top-down modulation of visual cortex processing. Science (80-) 345:660–665.

Zhou Y, Wang X (2014) Spatially extended forward suppression in primate auditory cortex. Eur J Neurosci 39:919–933.

